# FishPhyloMaker: An R package to generate phylogenies for ray-finned fishes

**DOI:** 10.1101/2021.05.07.442752

**Authors:** Gabriel Nakamura, Aline Richter, Bruno E. Soares

## Abstract

Phylogenies summarize evolutionary information that is essential in the investigation of ecological and evolutionary causes of diversity patterns. They allow investigating hypotheses from trait evolution to the relationship between evolutionary diversity and ecosystem functioning. However, obtaining a comprehensive phylogenetic hypothesis can be difficult for some groups, especially those with a high number of species, that is the case for fishes, particularly tropical ones. The lack of species in phylogenetic hypotheses, called Darwinian shortfalls, can hinder ecological and evolutionary studies involving this group. To tackle this problem, we developed FishPhyloMaker, an R package that facilitates the generation of phylogenetic trees through a reliable and reproducible procedure, even for a large number of species. The package adopts well-known rules of insertion based on cladistic hierarchy, allowing its use by specialists and non-specialists in fish systematics. We tested the reliability of our algorithm in maintaining important properties of phylogenetic distances running a sensitivity analysis. We also exemplified the use of the FishPhyloMaker package by constructing complete phylogenies for fishes inhabiting the four richest freshwater ecoregions of the world. Furthermore, we proposed a new method to calculate Darwinian shortfalls and mapped this information for the major freshwater drainages of the world. FishPhyloMaker will expand the range of evolutionary and ecological questions that can be addressed using ray-finned fishes as study models, mainly in the field of community phylogenetics, by providing an easy and reliable way to obtain comprehensive phylogenies. Further, FishPhyloMaker presents the potential to be extended to other taxonomic groups that suffer from the same difficulty in the obtention of comprehensive phylogenetic hypothesis.

**Highlights:**

- We provide the first automated procedure to check species names, construct phylogenetic trees and calculate Darwinian shortfalls for ray-finned fishes (Actinopterygii) by the R package FishPhyloMaker.
- This package provides functions to assemble phylogenies through a fast, reliable, and reproducible method, allowing its use and replicability by specialists and non-specialists in fish systematics.
- The package also provides an interactive procedure that gives more flexibility to the user when compared with other existing tools that construct phylogenetic trees for other highly speciose groups.
- The package includes a new method to compute Darwinian shortfalls for ray-finned fishes, but the rationale of the provided algorithm can be extended in future studies to be used in other groups of organisms

## Introduction

Phylogenies have been widely explored in ecology in the last decades due to the development of theoretical frameworks, numerical methods, and software (*e.g*., Webb et al. 2008; Felsenstein 1985). The research agenda in ecology and evolution encompasses phylogenetic approaches from organismal to macroecological-scale, including trait evolution, invasion ecology, metacommunity ecology, and ecosystem functioning (Cavender-Bares et al., 2009). Hence, comprehensive phylogenetic trees must be available to address those topics. Large phylogenies were primarily developed by combining source-trees and published-trees (the supertree approach), by concatenating different data matrices of systematic phylogenetic characters to generate a single tree (the supermatrix approach), or by a mix of both approaches (Haeseler, 2012; Smith et al., 2009).

Well-established phylogenies for most of the known species are available for some groups, such as terrestrial vertebrates (birds (Jetz et al., 2012), mammals (Upham et al., 2019), amphibians (Jetz and Pyron, 2018), squamates (Tonini et al., 2016), sharks (Stein et al., 2018), and plants (Magallón et al., 2015), which also have powerful tools to generate phylogenetic trees for local/regional pools of species (*e.g*., Webb & Donoghue 2005 for mammals and plants; Jin & Qian 2019 for plants, to the others see http://vertlife.org/phylosubsets/). Inversely, available phylogenies for bony fishes (Betancur et al., 2017; Rabosky et al., 2018) display issues related to the taxonomic position of some clades (e.g., non-monophyletic groups) and the lack of species representativeness. The latter issue hampers answering some questions on the ecology and evolution of ray-finned fishes by generating inaccuracy in estimates of phylogenetic signal, trait evolution, and phylogenetic diversity (Seger et al., 2013; Boettiger et al., 2012a), or even impeding their calculation.

Ray-finned fishes (Actinopterygii) exhibit a complex evolutionary history and high ecological diversity (Albert et al., 2020), making them an interesting group to address questions in the interface of ecology and evolution (*e.g*., Roa-Fuentes et al. 2019; Nakamura et al. 2020). The difficulty in obtaining phylogenetic information can hinder our efforts to understand fish ecology and evolution. Additionally, the lack of phylogenetic information for species, *i.e*., Darwinian shortfalls, is currently investigated in a few lineages (*e.g*., Freitas et al., 2021), which impedes the mapping of the relative demand of additional efforts needed in entire regions or clades to uncover the phylogenetic history of fishes. This problem urges a rapid solution in the context of the accelerated loss of species (Chase et al., 2020).

A short-term solution to tackle the Darwinian shortfall for ray-finned fishes would be coupling the phylogenetic information with cladistic classification to produce comprehensive phylogenies (Diniz-Filho et al., 2013). This solution is laborious and lacks reproducibility when adding many species, and the specific steps are not precisely documented when did “by hand” procedures (Webb et al., 2008). An alternative would be using molecular techniques to generate comprehensive phylogenies (e.g. Pie et al., 2021). However, it demands high expertise and high financial investment (Roquet et al., 2013), limiting factors for several institutions. Therefore, automatizing the procedures of constructing comprehensive phylogenies using the information from cladistic hierarchy, as suggested by Diniz-Filho et al (2013), provides a more reliable, accessible, and short-term solution for evolutionary ecologists. The technique produces reliable phylogenetic information for community phylogenetics (Li et al., 2019).

In order to tackle the problem of obtention of comprehensive fish phylogenies in a reliable and reproducible way, we developed the FishPhyloMaker. This freely available R package facilitates the obtention of phylogenetic trees for ray-finned fishes. FishPhyloMaker automates the insertion procedure of species in the most comprehensive phylogeny (Rabosky et al., 2018) of ray-finned fishes following their taxonomic hierarchy. We illustrated how the FishPhyloMaker package solves the problem of obtaining comprehensive phylogenies by constructing phylogenetic trees for species inhabiting more than 3000 freshwater basins globally (Tedesco et al., 2017). Further, we developed a new method to quantify the Darwinian shortfalls, which we illustrate by mapping the Darwinian shortfalls for the abovementioned basins. Finally, we performed a sensitivity analysis to evaluate how our method preserves characteristics of the phylogenetic tree (pairwise distances among species and evolutionary distinctiveness), even with a varying number of inserted taxa. Our package overcomes the main problems associated with manually building phylogenies for ray-finned fishes by following a specific and documented procedure and reducing the manual labor in large phylogenies.

## Methods

### Inside the Fish(PhyloMaker): an overview of the package

A stable version of FishPhyloMaker can be downloaded from the CRAN repository (https://cran.r-project.org/web/packages/FishPhyloMaker/index.html), and a development version is available at the GitHub repository (https://github.com/GabrielNakamura/FishPhyloMaker). All analyses shown here were performed using the development version of FishPhyloMaker.

FishPhyloMaker is a freely available R package containing three main functions, *FishTaxaMaker*, *FishPhyloMaker*, and *PD_deficit*. Below, we describe the functions to generate phylogenetic trees, highlighting the input data, intermediate steps, and output objects. Brief descriptions of the package functions are available in Table 1.

**Table 1:**
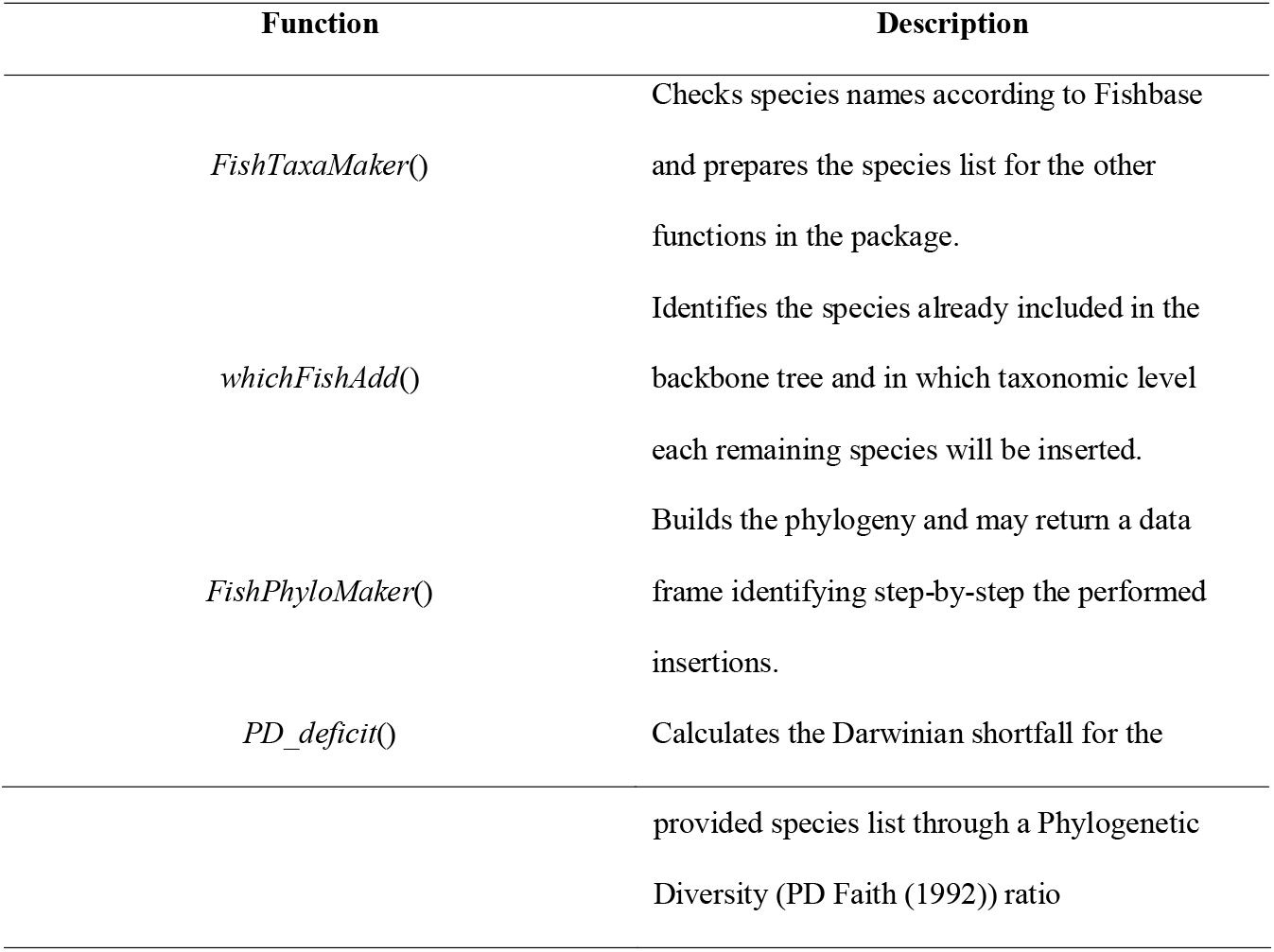
Functions presented in the package FishPhyloMaker and their descriptions.

### Fish TaxaMaker

The *FishTaxaMaker* function checks the validity of species names provided by the user and prepares a formatted data frame for the *FishPhyloMaker* function. The input data must be a string vector or a data frame containing a list of species from the regional pool or an occurrence matrix (sites x species). The genus and specific epithet (or subspecies) must be separated by underline (e.g., *Genus_epithet*). The function first classifies the provided species names as valid or synonymies based on Fishbase (Froese & Pauly, 2000) using the *rfishbase* package (Boettiger et al., 2012b). A new column summarizes names initially valid and the current valid names substituting identified synonymies. Unknown species to Fishbase are printed in the command line, and the user must manually inform the Family of these species. If the user types a Family not recognized in the FishBase, the user is asked to check the spelling and type the Order of this family. The output of the function is a list containing three elements: 1) a data frame displaying the taxonomic information (Valid name, Subfamily, Family, Order, Class, and SuperClass) for each species; 2) a data frame displaying the taxonomic information (Species, Family, and Order), only for the valid species; 3) a character vector displaying the species names not found in Fishbase.

### FishPhyloMaker

*FishPhyloMaker* is the core function of the package. This function builds a phylogenetic tree for the provided species list by inserting in and pruning species from the Rabosky et al., (2018) phylogenetic tree (Figure 1) downloaded by the fishtreeoflife R package (Chang et al. 2019). This phylogeny is the most up-to-date and comprehensive phylogenetic hypothesis for ray-finned fishes.

**Figure 1:**
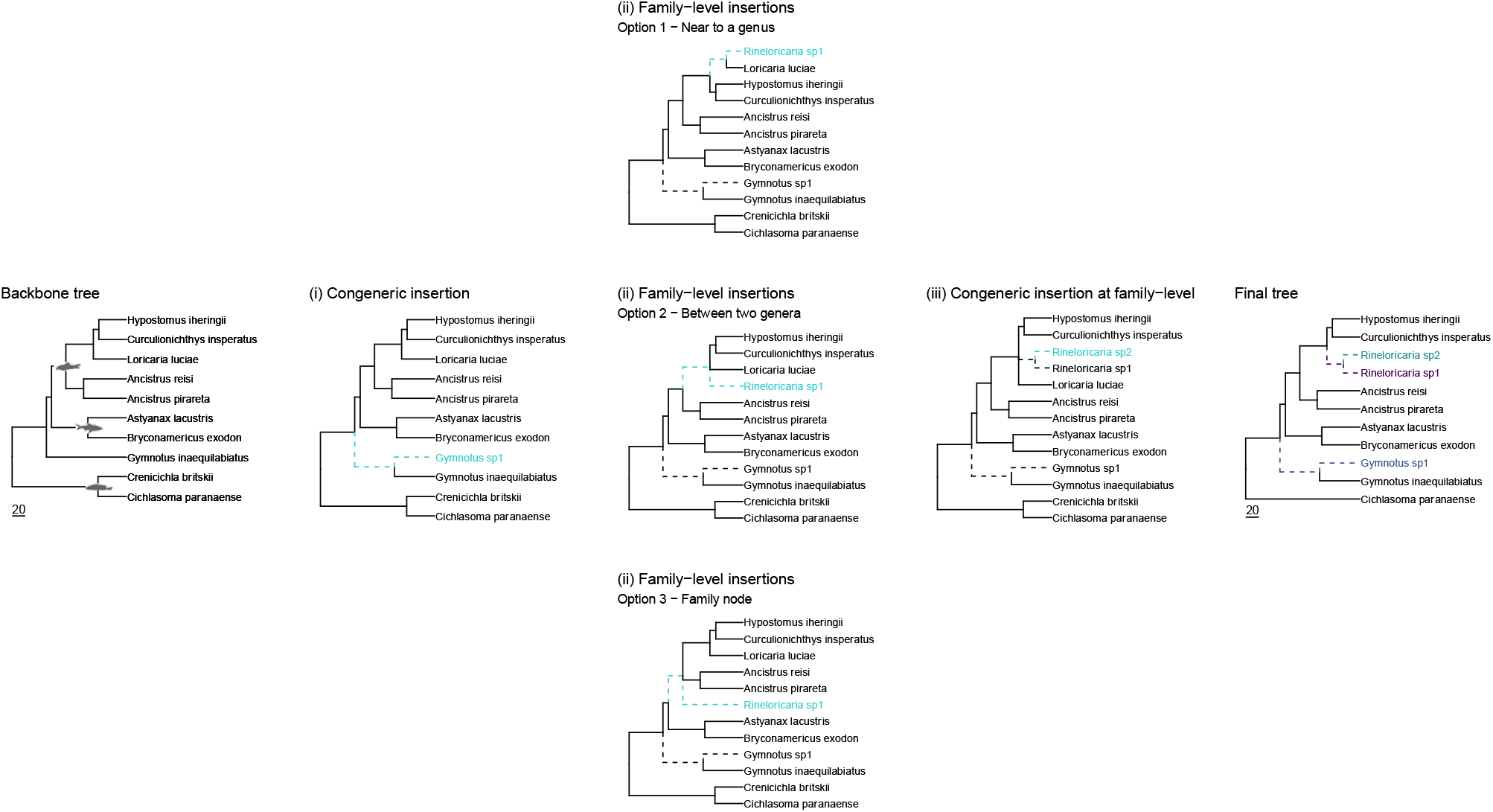
Schematic representation of insertion and subsetting procedure performed by the FishPhyloMaker() function. Here we used a hypothetical phylogeny containing ten species and four families (silhouettes inside the tree) as the backbone phylogeny. Step (i) represents the congeneric level of insertion. Step (ii) represents the three options that the user may choose in the Family-level round of insertions (Option 1 – near to a genus; Option 2 – between two genera; Option 3 – at the family node). (iii) represents the congeneric insertions at the family level and, finally, the final pruned tree containing only the species of interest.

The input for the *FishPhyloMaker* function can be the second element in the list returned by the *FishTaxaMaker* (Taxon_data_FishPhyloMaker) function or a manually constructed data frame with the same configuration (species, family, and order names for each taxon). The function also contains three logical arguments: insert.base.node, return.insertions and progress.bar. These three arguments are set by default as FALSE, TRUE, and TRUE, respectively, and allow the user to choose if the species must be at the base node of families/orders, if the insertions made by each species must be shown in the output and if a progress bar must be shown in the console.

The function works sequentially, first identifying which of the provided species are in the backbone phylogenetic tree (Rabosky et al., 2018). If all of them are already present in the backbone tree, the function returns a pruned one. If any of the provided species is not in the backbone tree, the function performs a four-level insertion routine. First, species from genera already included in the backbone tree are inserted as polytomies at the most recent ancestral node that links all congeneric species or as the sister species of the only species representing a genus in the backbone tree, as shown in *i* in Figure 1. In the case of *i* in Figure 1 the branch length is divided at half of its length and the species is inserted. Second, species not inserted in the previous step are then inserted at the family level by an interactive procedure using a returned list of all the genera within the same family of the target species. The user has the option to insert the target species as a sister taxon to a genus (*ii* in Figure 1, option 1, near to *Loricaria* genus), between two genera (*ii* in Figure 1, option 2, between genus *Loricaria* and *Hypostomus*), or at the node of the family (*ii* in Figure 1, option 3). If the user enters a single genus from the list, the function splits its branch and inserts the target as a sister taxon of this genus (option 1). If the user enters two genera separated by a blank space, the function inserts the target species as a polytomy at the most recent node that links the selected genera (option 2). If the user enters the family name, the function attaches the target species at the family node as a polytomy (option 3). Third, if any remaining species can now be inserted at the genus level, the function repeats the first procedure but records it as a Congeneric family-level insertion by splitting the branch length of the congeneric species at half of its length (*iii* in Figure 1). Fourth, remnant species are inserted at the order level following similar to the second step, by an interactive procedure using a returned list of all the families within the order of the target species. Hence, the user may specify a family to insert the target species as sister taxon (option 1), two families to insert it as a polytomy at the most recent node linking them (option 2), or the order to insert it as a sister taxon (option 3). The function will not perform insertions steps beyond the order level because it would add too much uncertainty to the phylogenetic tree.

Setting the argument `insert.base.node` as TRUE automatically inserts the target species from the second and fourth steps in the family and order nodes, respectively. This option facilitates the insertion of a large number of species or species with the unknown phylogenetic position. The default output is a list with two objects: (i) the pruned tree including only the provided species list (Final tree in Figure 1); (ii) a data frame identifying if each provided species was initially present in the backbone tree, in which step it was inserted, or not inserted at all. This data frame will flag each species with one of the six classification based on the insertion procedure: 1 – Present in tree will indicate species that were already present in the backbone tree; 2 – Congeneric insertion will indicate species that present at least one species of the same genus in the backbone tree and was inserted as congeneric of this species; 3 – Family insertion will indicate inserted species that did not present any congeneric species at backbone tree, but had at least one species of the same family in backbone tree; 4 – Congeneric at Family-level will indicate species that was added as congeneric after another species of the same genus was inserted at the Family level; 5 – Order insertion will indicate inserted species that did not presented any species of the same family in the backbone tree and must be inserted near to an extant family or in node corresponding to the order root in the backbone tree; 6 – not inserted will indicate species that did not present any species of the same order in the backbone tree, therefore was not inserted due their high uncertainty in the phylogenetic position.

### PD_deficit

The *PD_deficit* function calculates a measure of Darwinian shortfalls following Equation 1:

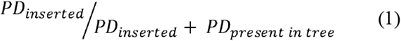

In this function, PD_inserted_ is the sum of the branch lengths of species in the phylogenetic tree before the insertion procedure. PD_present in tree_ is the sum of branch lengths of the species inserted in the tree. Therefore, the Darwinian deficit ranges from 0 (all species already present in the backbone tree before the insertion procedure) to 1 (all the species in the phylogenetic tree were inserted and were not presented in backbone phylogeny). PD_deficit function returns a vector with three values, the Darwinian shortfall (Equation 1), the total phylogenetic diversity calculated as the sum of branch lengths of the tree (*PD_total_*) with all species provided by the user, the sum of branch lengths inserted (*PD_inserted_*) in the tree and that was already present in the backbone tree (*PD_present in tree_*). It is worth noting that the sum of *PD_inserted_* and *PD_present_* are complementary, summing up to *PD_total_*. To calculate the Darwinian shortfall through the *PD_deficit* function, the user must provide a phylogenetic tree and a table of insertions, both obtained from the *FishPhyloMaker* function.

### Sensitivity analysis

We performed a sensitivity analysis to assess how the insertion procedure implemented herein and the amount of inserted species affect two characteristics of phylogenetic trees: the mean pairwise distance among species and the phylogenetic distinctiveness.

We 1) randomly change the name of a subsample of species within Rabosky’s phylogeny. Then, 2) we built a phylogeny for the species sampled with changed names in the previous step using the FishPhyloMaker function. Finally, we computed: 3) the matrix correlation (Pearson correlation) between the cophenetic distances of the subsampled species in Rabosky’s phylogeny and the FishPhyloMaker phylogeny; and 4) the Pearson correlation between the phylogenetic distinctness values for the Rabosky’s and FishPhyloMaker phylogenies. The evolutionary distinctness was calculated as the equal splits measure that is the sum of the contribution of all branches of a given lineage divided among its daughter branches (Redding and Mooers, 2006). Evolutionary distinctness measure was calculated using the phyloregion package (Daru et al., 2020).

The abovementioned steps (1, 2 and 3) were repeated 100 times for eleven different quantities (10%, 15%, 20%, 25%, 30%, 35%, 40%, 50%, 55%, 60%) of subsampled species from Rabosky’s phylogeny and inserted by the FishPhyloMaker function.

### Illustrating the use of FishPhyloMaker package

We provide an example of the usage of the *FishPhyloMaker* package by creating a phylogenetic tree using a global dataset of freshwater fishes inhabiting 3,119 freshwater drainage basins that cover more than 80% of the Earth surface and 14886 species (Tedesco et al., 2017). This dataset allowed in-depth investigation on the global patterns of species distribution and their evolutionary determinants (*e.g*., Miller & Román-Palácios, 2021). We built a phylogenetic for all species presented in Tedesco’s et al. dataset and mapped all the insertions realized. Moreover, we used this same dataset to demonstrate how to map Darwinian shortfalls, calculated following Equation 1 through *PD_deficit* function for all the drainage basins in the Tedesco et al. (2017) dataset. All the analyses were performed using the development version of the FishPhyloMaker package, which can be downloaded using the following command line:

~~~
devtools::install_github(“GabrielNakamllra/FishPhyloMaker”, ref = “main”, biiild_vignettes = TRUE)
~~~

We recommend that the user updates all the requested packages to avoid errors related to packages versions. We first prepared the fish occurrence by checking the validity of its names by using the function *FishTaxaMaker*. The occurrence matrix encompassed 14,886 species, from which 13,992 were valid names. The remaining 961 names were substituted by their corresponding valid names according to FishBase. We applied the *FishPhyloMaker* function to build a phylogenetic tree containing all the 14,886 species with valid names retrieved from *FishTaxaMaker* (Figure 2). For simplicity and reproducibility, we set the argument `insert.base.node` as TRUE, thus, inserting all species at the base node of its corresponding family and order when needed. We also set the argument `return.insertions = TRUE` for retrieving the insertion information of each species. Then, we applied the *PD_deficit* function to calculate the Darwinian shortfall for all the freshwater basins of the world harboring at least two species (Tedesco et al. 2017). The *PD_deficit* function was calculated considering congeneric insertions and insertions at the family level, however, the function may also include other levels of phylogenetic insertion, like order insertions. All the codes need to fully reproduce these analyses are provided at the GitHub repository (GabrielNakamura/MS_FishPhyloMaker). For further explanations and examples illustrating the usage of functions in the FishPhyloMaker package, the user can assess the package website https://gabrielnakamura.github.io/FishPhyloMaker/index.html and see the Articles section.

**Figure 2:**
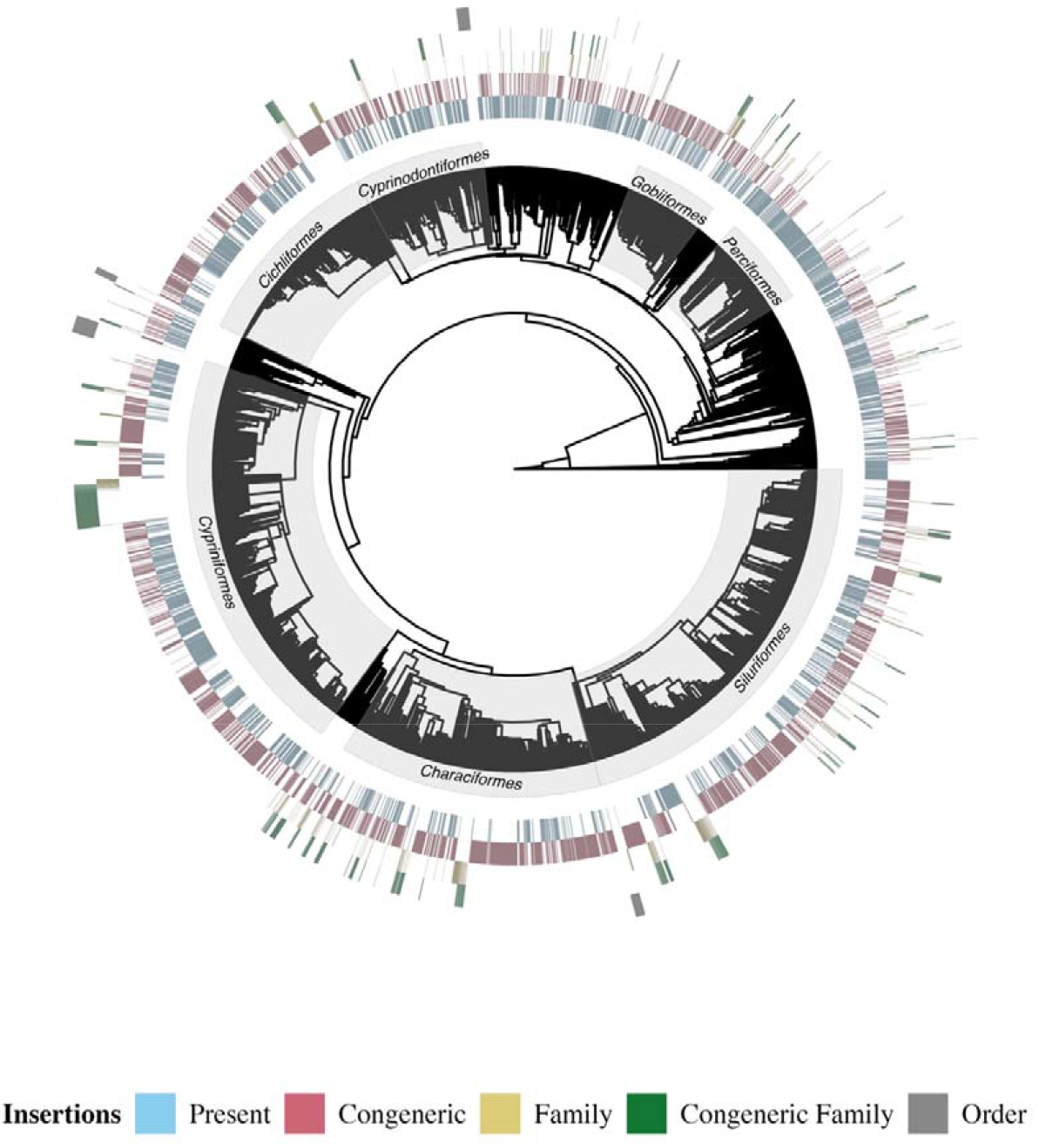
Phylogenetic tree obtained from FishPhyloMaker, containing 14,705 finned-ray species with their respective insertions. We also highlight in the gray rectangles the seven most speciose Orders.

## Results

The entire insertion procedure lasted approximately three hours using one core from a computer machine with an i5 processor. A total of 11,569 species were inserted, 6,418 species were already present in the backbone phylogeny, and 181 were not inserted at all, resulting in a phylogenetic tree containing 14,705 species (Figure 2). We also showed in Figure 2 all the insertions realized through the FishPhyloMaker function and the seven orders of ray-finned fishes with that present the highest number of species. We can see in Figure two that the insertions are evenly distributed throughout the phylogenetic tree.

We also depicted all the insertions made by FishPhyloMaker for all freshwater Ecoregion of the world. This was only possible because FishPhyloMaker flags all the insertions made during the insertion procedure. Figure 3 shows that Neotropics and Afrotropics regions exhibited the largest number of species inserted. On the contrary, despite the great area and number of basins, the Nearctic Ecoregion presented the smallest percentage of insertions, most of them congeneric. All Ecoregions and the percentage of species insertions per level are shown in Figure 3.

**Figure 3:**
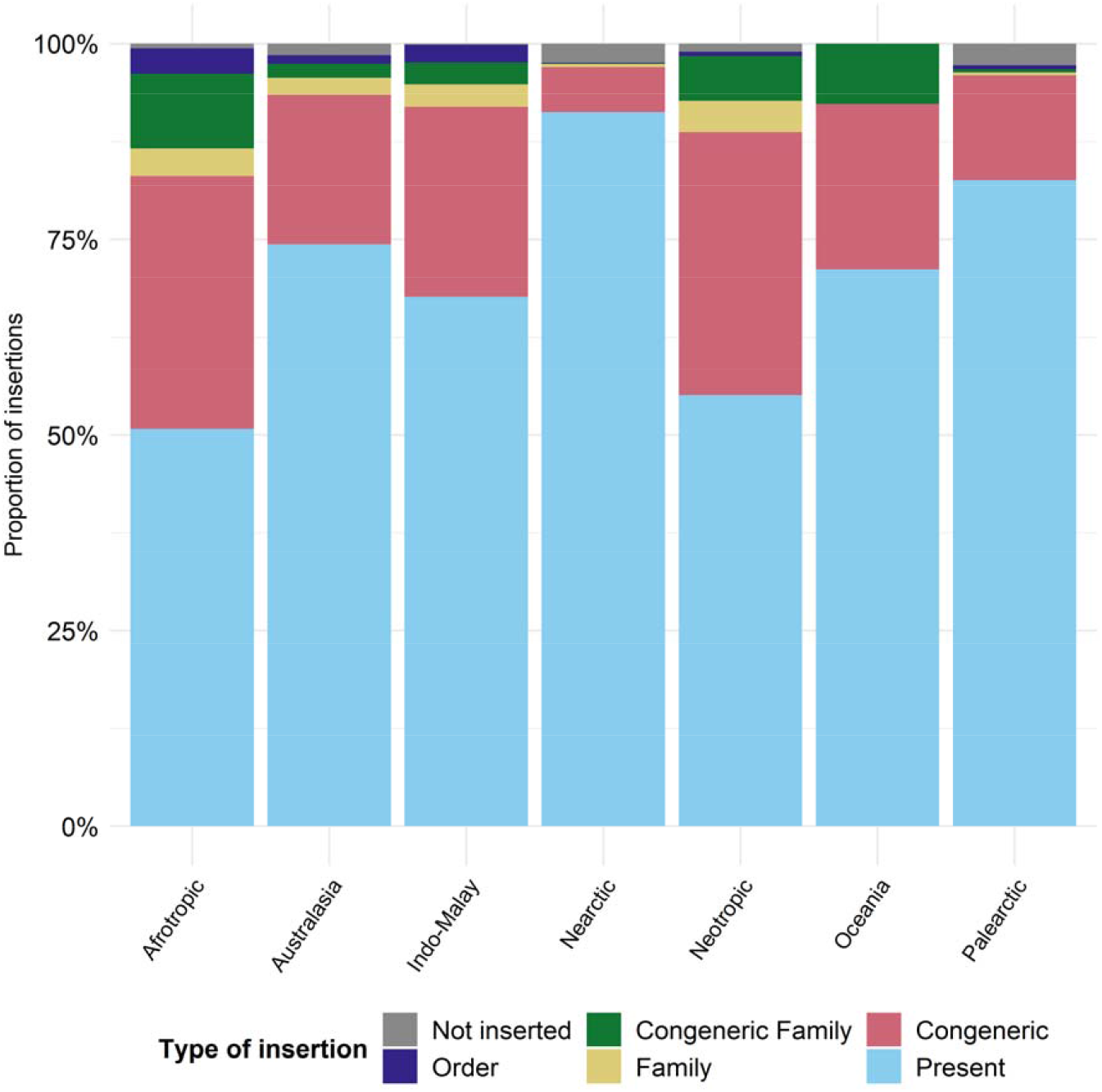
Barplot showing the percentage of species inserted in each one of the seven freshwater ecoregions of the world and their respective type of insertions mapped by FishPhyloMaker package.

We spatialized the Darwinian shortfalls per basin and observed that tropical regions exhibited the highest shortfalls, while northern sites had the lowest (Figure 4). The highest values of Darwinian shortfalls were found in Afrotropics and Neotropics, as some drainages did not harbor any (or only a few) species in the Rabosky’s phylogeny. The grey areas correspond to sites that do not present species occurrences accordingly to Tedesco et al. (2017) or presented less than two occurrences for the Order considered. We also depicted the Darwinian shortfalls for the four major orders in terms of species richness (bottom maps in Figure 4). For all the groups, the highest values of Darwinian shortfalls were found in the neotropical region, except for Cypriniformes, the group responsible for the highest values of Darwinian shortfalls in the watersheds in Asia and some basins in North America.

**Figure 4:**
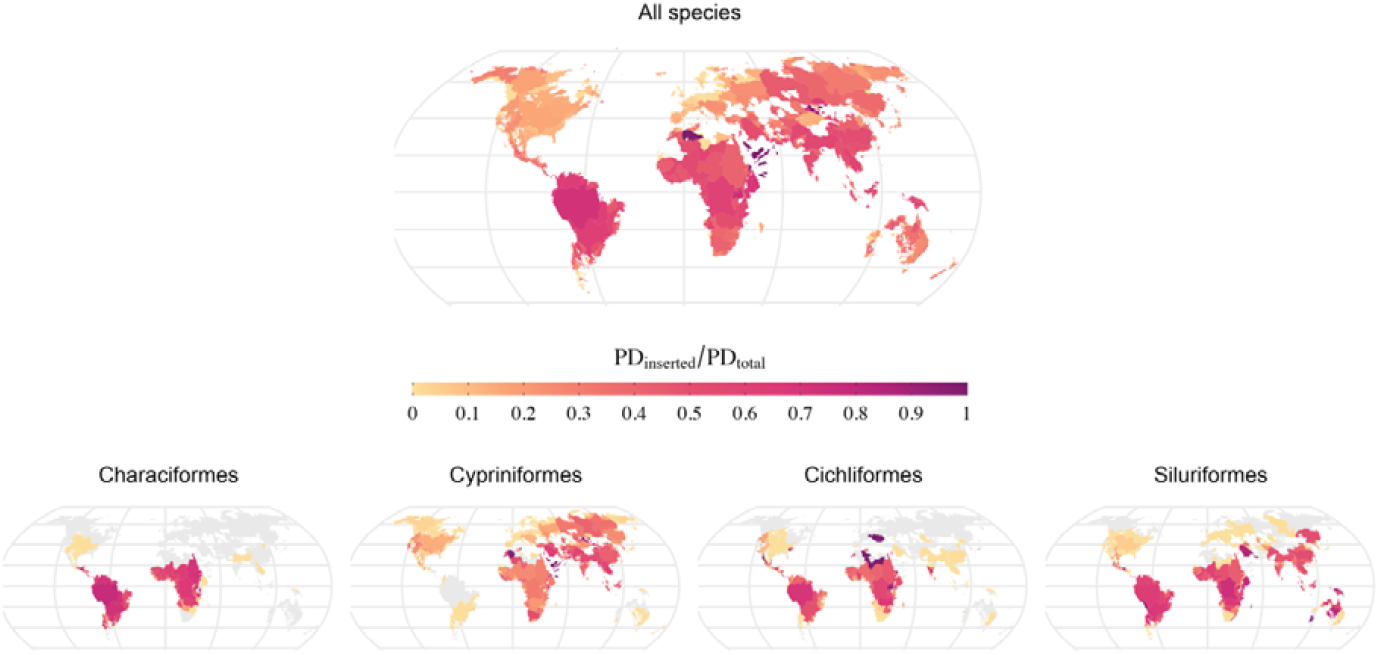
Global distribution of the Darwinian Shortfalls for ray-finned fishes, based on freshwater species occurrences in more than 3000 basins. Values near to 1 indicate a high Darwinian shortfall (a large number of congeneric insertions), while values near zero indicate low shortfalls. We depicted the Darwinian shortfall for the four major orders in terms of species richness (Characiformes, Cypriniformes, Cichliformes, and Siluriformes). Gray color indicates areas with no occurrence of species for a given order.

The sensitivity analysis highlights the strong correlation between the cophenetic distances of Rabosky’s and FishPhyloMaker phylogenies (Figure 5 B) even in varying levels of taxa insertions. Inversely, an increasing number of insertions on Rabosky’s phylogeny reduced the correlation between phylogenetic distinctness in the original phylogeny and that assembled by FishPhyloMaker (Figure 5 A).

**Figure 5:**
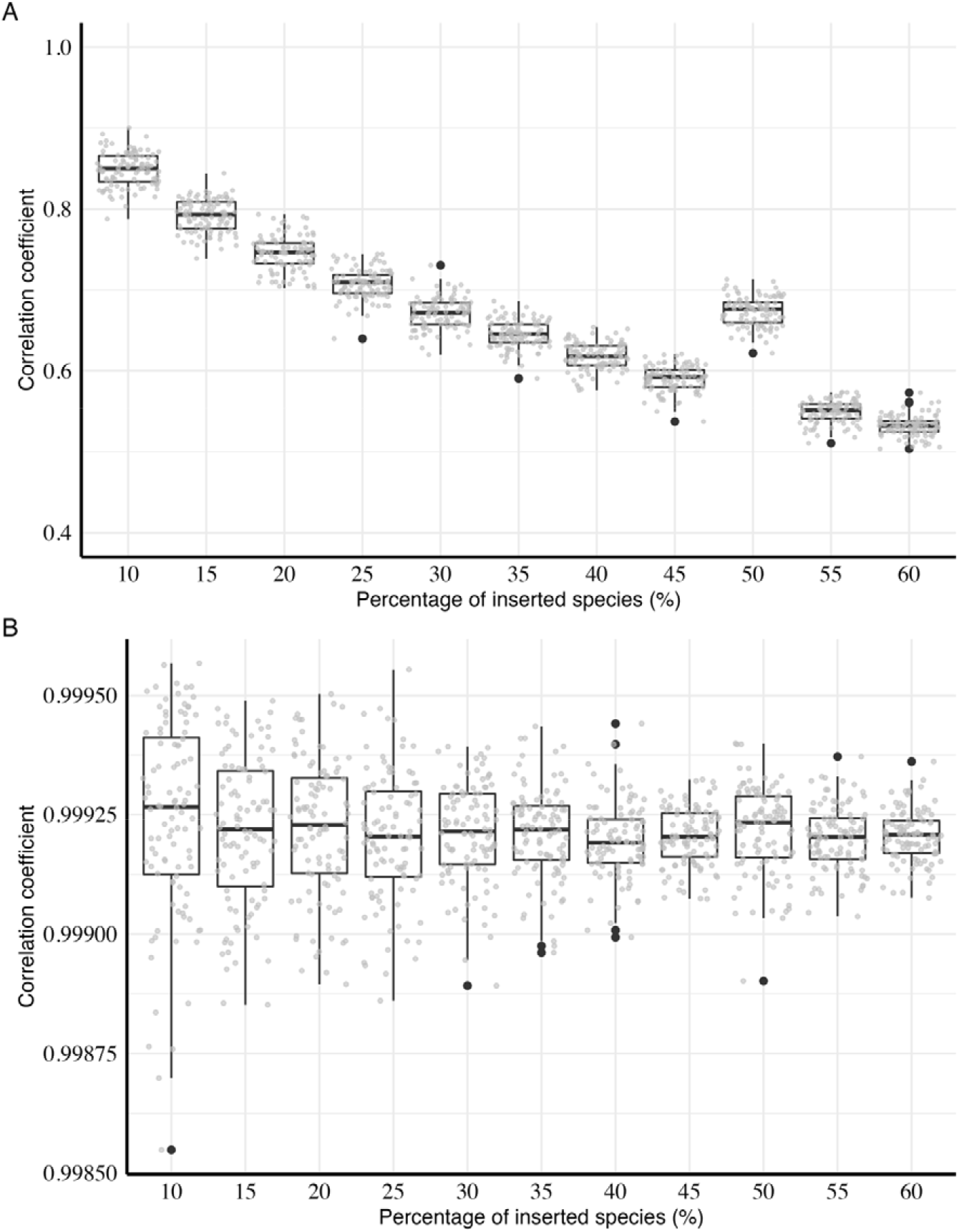
Barplots showing the correlation of evolutionary distinctness values (A) and between cophenetic distances (B) calculated from original phylogeny and inserted phylogenies with varying percentages of species inserted in the original phylogeny. Grey dots represent individual correlation values. The lower and upper hinges in boxplots represent the first and third quantiles while the middle hinge represents the median.

## Discussion

We provided a user-friendly, fast, reliable, and reproducible way to construct phylogenetic trees for a megadiverse group (Actinopterygii). The FishPhyloMaker package is in line with tools developed for plants, such as Phylomatic (C++ application) and V.PhyloMaker (R package) (Jin and Qian, 2019; Webb and Donoghue, 2005), but includes different features. These features include new options for inserting species through an interactive procedure in phylogenies and recording insertions. The latter feature allows a better systematization of building supertrees and calculating the first, to our knowledge, quantitative measure of the Darwinian shortfall.

Whereas Phylomatic allows the insertion of absent species only as congeneric or at the node corresponding to the family of the focal species (Webb and Donoghue, 2005), the FishPhyloMaker package delivers options through an interactive procedure of insertion. The performed insertions can be easily recorded in an R script, providing flexibility and the same level of reproducibility as other algorithms designed for similar purposes (*e.g*., Jin and Qian, 2019). This interactive option is a novelty when compared to similar insertion algorithms (*e.g*., Phylomatic).

The spatial distribution of the Darwinian shortfall is paramount to guide our future efforts to understand the history of life. The phylogenetic gaps in the knowledge of ray-finned fishes are geographically biased, with tropical basins presenting higher Darwinian shortfalls levels, as evidenced in this study. This gap in evolutionary knowledge could lead to a bias in evaluating the effects of evolutionary history and the interpretation of macroecological patterns for fish assemblages in these regions, which can affect conservation decisions based on the phylogenetic dimension of diversity (Assis, 2018).

Several biological and sociological factors can explain the observed bias in Darwinian shortfalls. First, the regions exhibiting the most significant Darwinian gaps also exhibit the largest freshwater fish diversity, which we can not describe at the same speed as less biologically rich areas (Hortal et al., 2015). Second, on-ground accessibility, human occupation, and economic development constrain investments in biodiversity research (Moura et al., 2018; Moura and Jetz, 2021), which is probably more pronounced in tropical regions than temperate ones, which may hamper field sampling and phylogenetic analyses.

Despite being more simple when compared with other insertion methods (e.g., Pearse and Purvis, 2013), FishPhyloMaker provided reliable results by preserving important characteristics of the phylogenetic tree, as we showed through the sensitivity analysis. Commonly used measures of phylogenetic diversity are based on the pairwise distance of species from a phylogenetic tree (e.g., Kraft et al., 2007; Webb et al., 2002), and we showed that the algorithm implemented in FishPhyloMaker successfully preserve the distances among species in the phylogenetic tree even for a great number of insertions.

### Limitations and possible applications

Future developments of the package should consider the Catalog of Fishes (van der Laan et al., 2021) to improve the nomenclature checking procedures. Despite Fishbase being a widely used database to check for the taxonomic classification of fishes, it may present delays in updating taxonomic information because it is not its primary purpose. Inversely, the Catalog of Fishes is an authoritative taxonomic list frequently updated.

An inherent limitation of the phylogenetic hypothesis produced by FishPhyloMaker is the large number of polytomies resulting from the insertion procedures. We recommend that users directly assess how the phylogenetic uncertainty affects further analysis when not using a fully solved phylogenetic tree (Martins et al., 2013). Furthermore, we recommend caution in the use of FishPhyloMaker phylogenies to compute measures that depend on speciation events (e.g., evolutionary distinctiveness and other split-based metrics) since the insertion procedure modifies the split events in the tree as shown in the sensitivity analysis.

These limitations do not preclude the package applicability for studies in phylogenetic community ecology since synthesis phylogenies do not significantly impact phylogenetic diversity indices as showed by previous studies (Li et al., 2019) and confirmed in ours (through sensitivity analysis). Moreover, this is the only automated tool able to provide a complete phylogenetic tree that can easily handle large datasets. FishPhyloMaker can be relevant for addressing several critical questions in ecology and evolution by facilitating the obtention of phylogenetic hypotheses for local pools of ray-finned fishes. This facilitation can be essential for regions with a large gap in the phylogenetic knowledge of fishes, such as the Neotropical region (Albert et al., 2020). Such phylogenetic hypotheses allow understanding how ecological traits evolved or how the current and past environmental conditions selected the lineages in different areas.

Biogeographical studies are usually restricted to one or a few lineages at larger scales due to the availability of molecular phylogenies (e.g. García-Andrade et al., 2021) or with phylogenies with a considerable number of absent species (Miller, 2021). The FishPhyloMaker package facilitates large-scale investigations on the biogeographic history of the most diverse group of vertebrates on Earth, the Actinopterygians, helping us understand the processes that drive this high diversity. Finally, we can map where the lack of phylogenetic information is the most critical once the function returns the insertion level of species. This information can directly elucidate the patterns of the Darwinian shortfalls for ray-finned fishes, contributing not only to direct sampling and studying efforts but also to evidence the need for increased efforts to decolonize science (Trisos et al., 2021). Therefore, we expect that the FishPhyloMaker package reduces the gaps and barriers to addressing ecological and evolutionary questions due to the difficulty or lack of a reliable phylogenetic hypothesis for local and regional pools of ray-finned fishes.

## Contributions

GN Conceptualization; Data curation; Formal Analysis; Methodology; Software; Writing – original draft. AR Data curation; Methodology; Software, Writing – review, and editing. BES Writing – original draft; Methodology.

## Acknowledgments

GN is a member of the National Institutes for Science and Technology (INCT) in Ecology, Evolution, and Biodiversity Conservation, supported by MCTIC/CNPq (proc. 465610/2014-5). BES and AR are grateful to FAPERJ and CAPES for their postdoctoral and doctoral grants, respectively. The authors also thank valuable suggestions made by LDS Duarte and other two anonymous Reviewers. Brazilian science resists.

